# Selectively Advantageous Instability and Information Theory in Sex-specific Aging

**DOI:** 10.64898/2026.09.02.748817

**Authors:** John Tower

## Abstract

Biological information is generally thought to be subject to selection for faithful maintenance. However, accurate preservation is often combined with regulated mechanisms that generate state change. Selectively advantageous instability (SAI) of biological information is modeled here as instability that is favored because useful alternatives become accessible. Modeling shows that active destabilization is favored above a threshold determined by environmental change, passive error, destabilization cost, and the relative adaptive targeting of active versus passive variation. When actively generated variation is sufficiently structured, selection simultaneously favors increased maintenance and increased active destabilization of the same information channel. This relationship is described as stabilization-destabilization complementarity. Shannon entropy quantifies uncertainty, whereas relative entropy quantifies mismatch between generated and fitness-relevant state distributions. Aging is not produced by reversible state-space exploration alone. Aging results when selected exploration also causes persistent or cumulative loss of maintained organization. Age and sex extensions show how delayed costs and shared genetic control can generate antagonistic pleiotropy and sexual conflict. A unified interpretation is provided in which SAI can be selected as a mechanism of adaptive state space exploration even when long term information displacement is costly.

## 1. Introduction

Biological inheritance and somatic function depend on the maintenance of information. DNA replication and repair, chromatin maintenance, proteostasis, and developmental canalization all help maintain established biological states. In terms of information-transmission these mechanisms increase fidelity between successive states. However, biological systems also contain mechanisms that function to generate change, including turnover, recombination, phenotypic switching, regulated mutagenesis, evolutionary capacitance, immune diversification, and extensive chromatin remodeling. The coexistence of high fidelity and active variation suggests that stability and instability need not occupy opposite ends of a single evolutionary continuum [1–3].

Selectively advantageous instability (SAI) is defined as instability of a replicator subunit that increases the replicative fitness of the replicator or the replicator lineage [3,4]. SAI is hypothesized to be an emergent property of replicators with multiple subunits and to be essential for life. SAI is proposed to include three basic mechanisms. The first is beneficial effects of free subunits, the second is adaptation to the environment and maintenance of genetic diversity, and the third is replacement of damaged subunits [5,6]. Consistent with these ideas, regulated instability and turnover of subunits is required for normal cell and organism function [3,7,8].

SAI is explored here in terms of information transmission and information theory [9]. Support for advantageous imperfect fidelity has been provided by studies of mutation rate modifiers, stress induced mutagenesis, evolutionary capacitance, and stochastic switching in fluctuating environments [1,10–13]. In the present study, a broader information theory formulation is developed in which passive information loss is separated from regulated destabilization and in which opposite selection on these two sources of uncertainty can be examined within the same biological information channel.

Here passive error is distinguished from active destabilization. Passive error is defined as residual corruption of biological information despite maintenance. Active destabilization is defined as a regulated process that increases transitions away from a currently maintained state. If both processes generate identical distributions of outcomes, paying to suppress one while paying to recreate the other is redundant. The key biological possibility is that active and passive variation may sample state space differently. Active destabilization may be more structured, conditional, or targeted toward historically useful alternatives. Once this distinction is introduced, high fidelity and active variation can become evolutionary complements rather than substitutes.

A minimal model is developed to establish four results. The central result is termed stabilization-destabilization complementarity. Sufficiently structured active variation can cause selection to favor greater passive fidelity while simultaneously maintaining a positive and costly active-destabilization rate in the same information channel. This separates the amount of uncertainty from its mechanistic source and state-space distribution. The invasion threshold for active destabilization is derived and shows that passive error can approach zero while optimal active destabilization remains positive. Finally, the framework is extended across age and sex to antagonistic pleiotropy, evolution of aging, and sexual antagonistic pleiotropy.

The aging interpretation is separated from the entropy result. Increased state entropy is treated as a consequence of selected state generation, whereas aging is defined by progressive information displacement that is accompanied by declining biological function. This distinction allows selected instability to be connected to antagonistic pleiotropy without requiring every entropy increasing process to cause aging.

Two complementary constructions are presented involving standard Shannon quantities. In a minimal two-subunit AB replicator model, selective degradation is shown to be favored despite an immediate functional cost. In the cellular factor turnover model, reversible degradation and synthesis are shown to generate cellular state space exploration. These constructions are then generalized with a transition kernel in which passive error and structured active destabilization are separated.

## 2. Materials and Methods

Mathematical calculations, simulations, figure calculations and generation of figures were performed using ChatGPT (GPT-5.6 Sol; OpenAI, San Francisco, CA, USA). Several mathematical calculations, simulations and figure calculations were also analyzed using Academia Co-scientist (model ID claude-opus-5; San Francisco, CA, US). All resulting calculations were reviewed, verified and interpreted by the author. Exact sample sizes and test statistics are reported in Results. A reproducibility archive is provided as supplementary material (SAI_reproducibility_archive_V23.zip). Reproduction of the computational analyses is coordinated by the master script reproduce_all.py. The theoretical figures are regenerated from the equations and parameter values reported in the manuscript. Software requirements and a manuscript to code output manifest are included in the archive. Additional details of materials and methods are provided in the Supplementary Materials.

## 3. Results

Symbols are defined again immediately below the equation in which they first appear. Distinct symbols are used across the principal submodels to avoid changes of meaning across sections, and are summarized in Supplementary Materials (Supplementary Table 1).

### 3.1 Minimal replicator model of selected instability and Shannon state entropy

A minimal molecular replicator was first considered [3,4] so that the selective and information theoretic consequences of instability could be separated from the more general transition model developed below. The replicator was assumed to contain two subunits A and B. The intact complex AB was assumed to replicate from substrate, and state specific replication rates were assigned to A, AB, and AAB. The A subunit was also allowed to associate with AB to form AAB. Free A was assumed not to replicate, whereas AAB was assumed to replicate more effectively than AB. Thus *R*_A_ = 0 and *R*_AAB_ > *R*_AB_ > 0.

Differential subunit stability was then introduced. The B subunit was assumed to be less stable than A. Loss of B from AB therefore generated free A according to

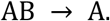

> Here AB is the intact two subunit replicator, A is the free A subunit produced by loss of B, B is the less stable subunit, and → denotes the transition produced by B degradation.

This transition imposed an immediate cost because a competent AB replicator was destroyed and the resulting A state was unable to replicate. The same transition also generated the A required for formation of the more effective AAB replicator according to

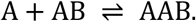

> Here + denotes association of molecular species, AAB is the three subunit replicator containing two A subunits and one B subunit, and ⇌ denotes a reversible association and dissociation process.

The selective effect of B instability was therefore determined by a competition between loss of AB and access to AAB. Let d_B_ denote the B destabilization rate, γ the effective return rate from A to AB, u the loss rate from AAB, and χ(d_B_) the rate at which AAB becomes accessible as destabilization generates A. For small d_B_, χ(d_B_) can be written as k_1_ d_B_ plus terms of higher order.

At age or elapsed time t, the first order occupancies generated by a small positive d_B_ are

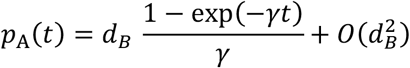

> Here p_A_(t) is the probability or first order occupancy of state A at elapsed time t, d_B_ is the B destabilization rate, γ is the effective return rate from A to AB, exp denotes the exponential function, and O(d_B_^2^) denotes terms of second and higher order in d_B_.

and

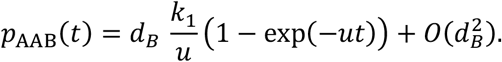

> Here p_AAB_(t) is the probability or first order occupancy of state AAB at elapsed time t, k_1_ is the first order coefficient relating access to AAB to d_B_, and u is the loss rate from AAB. The quantities d_B_, exp, and O(d_B_^2^) are defined above.

The remaining probability is assigned to AB to the same order. The instantaneous effect of destabilization on replication is therefore

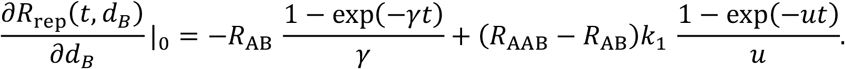

> Here R_rep_(t,d_B_) is the instantaneous replication output at time t and destabilization rate d_B_, ∂ denotes a partial derivative, the vertical bar with subscript 0 denotes evaluation at d_B_ equal to zero, R_AB_ is the replication rate of AB, and R_AAB_ is the replication rate of AAB. The remaining symbols are defined above.

Positive B instability can consequently be favored even though B degradation destroys a functional replicator. The loss is favored when the reproductively weighted gain produced through access to AAB exceeds the reproductively weighted cost produced through occupancy of A. This provides a minimal realization of SAI in which destruction of part of a functioning replicator is selected because the products of that destruction alter the accessible state space and increase lineage replication.

The same model yields a direct Shannon result. The state variable X was defined over the set containing AB, A, and AAB. In the corresponding no destabilization system, the maintained reference distribution is *P*_0_ = (1,0,0), for which *H*(*X*) = 0. For every positive d_B_ and every positive elapsed time, probability is transferred from AB into A and AAB. The distribution therefore becomes nondegenerate and

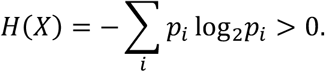

> Here H(X) is the Shannon entropy of state variable X, X takes the states AB, A, and AAB, Σ_i_ denotes summation over all states indexed by i, p_i_ is the probability of state i, log_2_ is the base 2 logarithm, and > denotes greater than.

For small d_B_, if 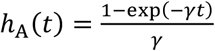 and 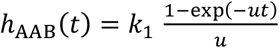, the leading entropy contribution is

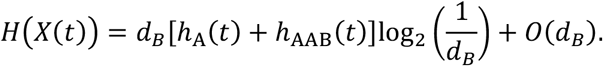

> Here X(t) is the state variable at elapsed time t, h_A_(t) is (1 − exp(−γ t)) divided by γ, h_AAB_(t) is k_1_(1 − exp(−u t)) divided by u, and O(d_B_) denotes terms whose magnitude is of order d_B_ as d_B_ approaches zero. The remaining symbols are defined above.

Therefore, when positive destabilization is selected in this minimal model, a system with greater Shannon state entropy than the corresponding no destabilization system is necessarily selected. Entropy is not assumed to be a target of selection and entropy maximization is not implied. Increased entropy is produced as a consequence of the selectively advantageous state generating process.

A distinction between state entropy and aging is required. In the minimal replicator model, Shannon quantities are defined for an ensemble distribution over AB, A, and AAB rather than for a single AB molecule. An individual AB replicator undergoes the discrete transition from AB to A when B is lost. Progressive information displacement is therefore an ensemble property. Shannon entropy need not increase monotonically with elapsed time because an ensemble displaced from the intact reference can later become concentrated in a different state. If *δ*(*t*) = 1 − *p*_AB_(*t*), then δ(t) increases to first order with positive d_B_. The Jensen Shannon divergence between the ensemble distribution at time t and the intact reference P_0_ = (1,0,0) therefore also increases to first order. More precisely, using base-2 logarithms, if δ is the total probability mass displaced from the degenerate AB reference state, the Jensen Shannon divergence approaches δ divided by 2 to first order, independently of how the displaced mass is partitioned between A and AAB.

At the level of an individual replicator, no additional functional variable is required to represent the immediate detrimental effect of destabilization. Functional integrity is represented by the intact replication competent AB state. Loss of B converts an individual AB replicator into nonreplicating A. Therefore each B degradation event imposes a local functional cost while also generating the A that permits formation of the superior AAB replicator elsewhere in the ensemble.

Opposing consequences can therefore be produced by the same selected instability at different levels. An individual replication competent AB is lost through conversion to A, whereas replication of the ensemble can be increased through formation of AAB. The detrimental effect is intrinsic to the destabilizing event and does not require a separate detrimental process.

A concrete biological interpretation is obtained when the replicator ensemble is placed within a bounded individual system such as a cell. The cell can contain many AB replicators whose states collectively define an intracellular distribution over AB, A, and AAB. Continued operation of B destabilization then allows the intracellular ensemble to sample states that would remain inaccessible under perfect preservation of AB. If the relative replication or functional value of these states depends on environmental conditions, this state space exploration can increase the ability of the cell and its replicator lineage to respond to environmental change.

The adaptive value of this exploration does not require Shannon entropy itself to be favored. Selection is instead applied to the transition process and to the fitness consequences of the states that become accessible. Perfect preservation restricts the ensemble to the existing AB state. Positive destabilization generates access to A and AAB, and more generally to alternative states in an extended system. When environmental conditions vary, a nonzero transition rate can therefore be favored because the expanded accessible state space increases the probability that a useful state is occupied. Increased Shannon state entropy can accompany this exploration without being its evolutionary objective.

The same state space exploration carries an intrinsic cost. Each transition from AB to A destroys a replication competent AB and produces a nonreplicating state. Some of the released A can contribute to formation of the superior AAB state, but the exploratory process necessarily includes loss of maintained AB organization. Repeated operation of the selected destabilization process can therefore produce both adaptive opportunities and cumulative displacement of the intracellular ensemble from its initially maintained distribution. The benefit of exploration and the cost of maintaining organization are generated by the same mechanism.

Aging can be mapped onto this tradeoff at the level of the cell or organism rather than the individual AB replicator. An individual AB undergoes a discrete transition and does not gradually age in the model. The cell can age as the distribution of its constituent informational units is progressively displaced from a maintained youthful distribution. The minimal model therefore establishes selected instability, local functional loss, state space exploration, and ensemble information displacement. The further claim that this displacement contributes to aging is conditional on cumulative displacement impairing cellular or organismal function. Under that condition, aging can represent a long term cost of an evolutionarily advantageous capacity to explore biological state space.

### 3.2 Reversible factor turnover as cellular state space exploration

A second construction was used to map SAI directly onto cellular states. A cell containing factor B was assigned the state B present, and a cell lacking B was assigned the state B absent. Degradation of B at rate k_d_ and synthesis of B at rate k_s_ were represented as reversible transitions between these states. At stationarity, the B present probability is k_s_ divided by k_s_ plus k_d_, and the B absent probability is k_d_ divided by k_s_ plus k_d_. A positive degradation rate therefore makes an alternative cellular state accessible, while return to the B present state is permitted by synthesis. Active information destabilization does not require synthesis to be impossible or imperfect. The physical state present at an earlier time can be lost through degradation even when the same coarse grained state can later be reconstructed by synthesis.

Two distinct Shannon consequences were considered. The Shannon state entropy H(X) quantifies uncertainty in the instantaneous distribution of B present and B absent states. The temporal mutual information I(X_t_; X_t+Δt_) quantifies how much information about an earlier cellular state is retained at a later time. State entropy can therefore remain constant while temporal memory is altered. Increased entropy is not required for active information destabilization when information about the past state is lost through faster turnover.

For the reversible two state process, memory of the initial B state decays with a characteristic correlation time *τ*_*c*_ = 1/(*k*_*d*_ + *k*_*s*_). If both rates are multiplied by the same positive factor, the stationary probabilities and stationary state entropy are unchanged, whereas the correlation time is shortened by the same factor. Information about the past state is therefore lost more rapidly even though the stationary distribution is unchanged. Reversible turnover can consequently be interpreted as controlled biological forgetting as well as cellular state space exploration.

#### 3.2.1 Turnover rate and temporal information can vary independently of state entropy

The distinction can be stated directly. If both turnover rates are multiplied by the same factor *λ* > 1,

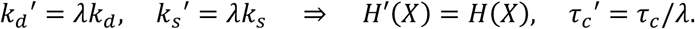

> Here k_d_ and k_s_ are the degradation and synthesis rates of factor B, respectively, the prime denotes the corresponding rate after proportional acceleration, λ is the positive multiplicative factor applied to both turnover rates, ⇒ denotes implication, H(X) is the stationary Shannon state entropy of state variable X, H′(X) is the stationary Shannon state entropy after proportional acceleration, τc is the temporal correlation time before acceleration, and τc′ is the temporal correlation time after acceleration.

Thus stationary state entropy is unchanged while temporal memory is shortened. For a fixed interval Δt, the state-dependence factor changes from

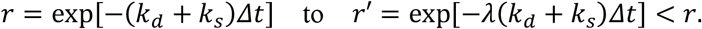

> Here r is the state dependence factor over elapsed interval Δt before proportional acceleration, r′ is the corresponding state dependence factor after acceleration, Δt is the elapsed time interval.

Consequently, for *λ* > 1, temporal mutual information *I*(*X*_*t*_; *X*_*t*+*Δt*_) decreases even though H(X) is unchanged. Reversible turnover can therefore alter temporal information retention independently of instantaneous state entropy.

The two cellular states were allowed to differ in fitness across environmental states. When B present is favored in one environment and B absent is favored in another, turnover can be selected because cellular states that match different future environments are sampled [3]. Perfect preservation of B maximizes retention of the previous B present state but prevents this route of exploration. SAI is obtained when the expected fitness gained through access to useful alternative states exceeds the costs imposed by degradation, replacement, and occupancy of mismatched states. Selection is therefore applied to the rate and structure of forgetting rather than to entropy itself.

The construction was generalized to *n* reversible factors F1 through F*n*. Each factor was assigned present and absent states. The cellular state at time *t* was represented by

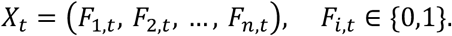

> Here Xt is the joint cellular state at time t, Fi,t is the state of factor i at time t, F denotes a reversible factor, i indexes the factors from 1 through n, n is the total number of reversible factors, 0 denotes the absent state, 1 denotes the present state, ∈ denotes membership in the set, braces denote the set of allowed binary factor states, and parentheses denote the joint vector of factor states.

If all combinations are accessible, the number of possible cellular states is

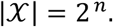

> Here *X* is the cellular state space, |*X*| is the cardinality of that state space, vertical bars denote cardinality, 2 is the number of possible states of each factor, n is the number of reversible factors defined above, and 2^n^ is the maximum number of joint cellular states when all binary combinations are accessible.

Degradation and synthesis rates determine both the distribution among accessible states and the rate at which information about previous states is lost. Adaptive value is obtained when selected turnover reduces dependence on obsolete past states while increasing access to states that are useful under recurrent future environmental conditions. Passive error and SAI can therefore produce similar instantaneous uncertainty while differing in temporal structure and fitness consequence.

### 3.3 Definition of a biological information channel

The term “biological information channel” is used to denote a biological process through which a current state X_t_ influences a subsequent state *X*_*t*+*Δt*_, mathematically represented by the conditional transition distribution *Q*(*X*_*t*+*Δt*_ | *X*_*t*_). The state X_t_ is the information state and the channel is the probabilistic transition rule connecting present and subsequent states [9]. Therefore passive error and active destabilization modify the channel, whereas their outcomes are changes in the information state. The term does not imply a particular physical medium or that biological information has semantic meaning in the Shannon sense. An individual organism is therefore not treated as a single information channel. Instead, it is more realistically represented as a network of coupled channels transmitting genomic, epigenetic, transcriptional, proteomic, metabolic, cellular, and physiological states through time.

#### 3.3.1 From factor turnover to temporal information loss and ontogenetic channels

The reversible factor model was used as a direct molecular realization of the general state transition framework. For a single factor B, two cellular states were made accessible by degradation and synthesis. The cellular state at time t can therefore be represented by X_t_, and the subsequent state can be described by the conditional transition distribution *Q*(*X*_*t*+*Δt*_ | *X*_*t*_). Physical degradation changes the present cellular state and can reduce information retained about its previous state even when synthesis later restores B.

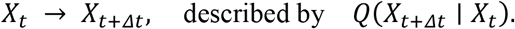

> Here X_t_ is the cellular information state at time t, Δt is the elapsed time interval, X_t+Δt_ is the state after that interval, Q(X_t+Δt_ | X_t_) is the conditional probability distribution of the later state given the earlier state, → denotes temporal transition, and the vertical bar denotes conditioning on the earlier state.

The same construction can be extended to n factors. When each factor F_i_ can be present or absent, the cellular state can be represented by *X*_*t*_ = (*F*_1,*t*_, *F*_2,*t*_, …, *F*_*n,t*_). As many as 2^*n*^ cellular states can thereby be made accessible. The degradation and synthesis rates of the factors determine transitions through this state space. Perfect stability restricts exploration and preserves information about previous factor states, whereas nonzero turnover permits alternative states to be sampled and permits temporal information about earlier configurations to decay.

Two information quantities were therefore separated. The state entropy H(X_t_) quantifies uncertainty in the distribution among cellular states at a specified time. The temporal mutual information *I*(*X*_*t*_; *X*_*t*+*Δt*_) quantifies information about the earlier state that remains in the later state. For the reversible B process, proportional acceleration of both k_d_ and k_s_ leaves the stationary state probabilities unchanged but shortens the correlation time. The same stationary state entropy can therefore be maintained while temporal mutual information is lost more rapidly.

Environmental matching can then be represented within the same construction. A cellular transition kernel *Q*(*X*_*t*+*Δt*_ | *X*_*t*_) is determined in part by molecular parameters such as *k*_*d,i*_ and *k*_*s,i*_, while a distribution over useful future states is supplied by the environmental process. Selection can therefore be applied to molecular transition rates according to the fitness consequences of retained memory and newly accessible states. Information about a past cellular configuration can be selectively discarded when reduced dependence on that configuration increases access to states that are useful under future conditions.

The resulting tradeoff can be expressed as a balance between retention of information about the past and exploration of states relevant to the future. Too little turnover can preserve obsolete cellular information and restrict adaptation after environmental change. Excessive turnover can erase useful memory and generate mismatched states. An intermediate rate of active destabilization can therefore be favored when the temporal statistics of the environment reward both persistence and switching.

The construction can be applied directly to ontogenetic change by allowing x to denote age rather than arbitrary time.

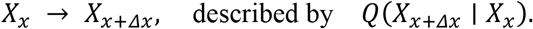

> Here X_x_ is the biological information state at age x, Δx is an age interval, X_x+Δx_ is the state after that interval, and Q(X_x+Δx_ | X_x_) is the conditional probability distribution of the later state given the state at age x.

A chromatin state, transcriptional program, mitochondrial state, cell identity, or multivariate physiological state can be represented by X. Declining temporal mutual information with age can describe loss of dependence on earlier biological states, while increasing conditional entropy can describe reduced predictability of later states. Neither quantity alone establishes molecular damage. Passive error, reduced maintenance, selected state change, or combinations of these processes can produce these information changes.

Reversible exploration and temporal forgetting alone are not defined as aging. Aging can nevertheless be produced even when the exploratory factor is reconstructed accurately. A transient state that is useful under a current or near future environment can produce a downstream consequence that persists after the exploratory factor has returned to its previous state. The factor state can therefore be restored while the larger cellular state remains altered. Aging is predicted when persistent consequences of repeatedly occupied adaptive states accumulate and progressively impair later function. Imperfect reconstruction and age dependent changes in transition rates remain possible mechanisms, but neither is required.

The ontogenetic channel must be distinguished from the intergenerational channel through which transmission from parents to descendants is described.

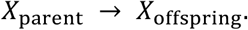

> Here X_parent_ is the heritable information state of the parent, X_offspring_ is the corresponding state transmitted to the offspring, and → denotes intergenerational transmission.

Natural selection is exerted through the intergenerational process. Alleles and other heritable determinants of within organism information dynamics are changed in frequency according to their effects on survival and reproduction. Shannon fidelity within an aging organism is therefore not directly maximized by selection. Heritable mechanisms controlling the fidelity and transition structure of within organism channels are favored only insofar as lifetime reproductive fitness is increased.

The two levels are linked causally. Ontogenetic information dynamics are determined by heritable regulatory architecture, survival and reproduction are influenced by those dynamics, and the frequency of the heritable architecture is changed by differential reproduction.

heritable regulation → within organism information dynamics → lifetime fitness → evolutionary change

> Here heritable regulation denotes genetically or otherwise heritably specified control of information dynamics, within organism information dynamics denotes state transitions during the lifetime, lifetime fitness denotes survival and reproductive output integrated across life, evolutionary change denotes change in frequencies of heritable determinants across generations, and → denotes causal influence in the proposed hierarchy.

A late life departure from a youthful state can therefore be deleterious at the level of the individual even when the mechanism that caused or permitted the transition was favored because an earlier reproductive, stress response, or plasticity benefit was generated. This is consistent with the antagonistic pleiotropy theory of aging [14].

#### 3.3.2 Scale-generality and nested biological information channels

This information-theoretic SAI framework is scale-general rather than tied to a particular biological level. Its minimal object is a state transition *X*_*t*_ → *X*_*t*+*Δt*_ governed by a transition kernel *Q*(*X*_*t*+*Δt*_ | *X*_*t*_), with maintenance suppressing passive departures, passive error generating comparatively unstructured variation, and active destabilization generating regulated variation. The biological interpretation of X, the relevant time step, and the fitness consequences of the transition must nevertheless be specified independently at each organizational level [2,3].

At the intergenerational level, X may denote a heritable genetic or epigenetic state. Maintenance includes replication fidelity and repair, and passive error includes spontaneous mutation. Candidate active-destabilization mechanisms include regulated mutagenesis, recombination, gene conversion, gene destruction, or other evolved processes that generate heritable variation. The framework predicts that high ordinary transmission fidelity can coexist with positively selected mechanisms that deliberately generate variation.

At the organismal or ontogenetic level, X_x_ may denote the somatic state of an individual at age x. The transition *Q*(*X*_*x*+*Δx*_ | *X*_*x*_) can describe maintenance and remodeling of chromatin, transcriptional programs, mitochondrial function, cell identity, proteostasis, metabolism, or physiology. Here the information dynamics occur within the individual, whereas evolutionary optimization occurs indirectly through the effects of heritable regulators of those dynamics on lifetime survival and reproduction.

At the cellular level, X may represent stem-cell, differentiated, stress-response, immune, senescent, or other cellular states. A cell can require high-fidelity preservation of some components while retaining regulated transitions in others. Stabilization to destabilization complementarity therefore predicts that cellular fidelity and cellular plasticity need not be opposing scalar traits.

At the molecular level, X can represent a DNA sequence, RNA or protein population, chromatin configuration, regulatory-network state, protein conformation, organelle state, or other molecular or subcellular state. Destabilization can include regulated physical degradation, including RNA decay, protein degradation, and selective destruction of organelles such as mitochondria. Such degradation can reduce persistence of the current state and thereby permit replacement, reconstruction, or exploration of alternative states. A molecular transition or degradation event should not itself be described as selectively advantageous merely because it is regulated. Rather, SAI requires that heritable regulatory architecture generating or permitting the destabilization has been favored because the resulting loss, turnover, replacement, or state variation increases reproductive fitness directly or through higher-level consequences.

At the population level, X may denote a distribution of phenotypes or states. Bet hedging and stochastic phenotypic switching provide natural examples in which evolved mechanisms generate a distribution of outcomes rather than maximally preserving a single state. These cases are closely related to existing information-theoretic treatments of adaptation in fluctuating environments [1,13,15].

The common architecture across scales can be summarized as

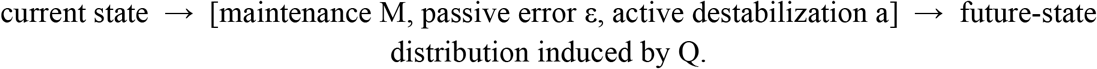

Here M is maintenance investment, ε is the passive error probability, a is the probability of active destabilization, Q is the biological transition kernel that generates the future-state distribution, brackets group the three processes acting on the current state, and → denotes the mapping from current state through these processes to the future state distribution.

A fitness-relevant environment or target process supplies a distribution P over useful future states. Selection can favor reduction of mismatch between P and the future-state distribution induced by Q, subject to the costs of maintenance, destabilization, and delayed consequences. The level of biological organization changes the interpretation of P and Q, but not the basic mathematical logic.

Biological information channels are also nested. Molecular transitions alter cellular transition probabilities, cellular transitions alter tissue and organismal physiology, organismal states alter survival and reproduction, and differential reproduction changes the frequencies of heritable regulators across generations.

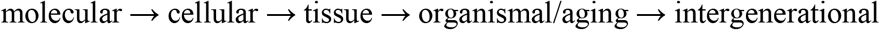

> Here molecular, cellular, tissue, organismal or aging, and intergenerational denote nested levels of biological organization, and → denotes propagation of state effects from one level to the next.

As a consequence, the site at which destabilization occurs need not be the level at which its selective advantage is realized. For example, an actively regulated chromatin transition may produce cellular plasticity, improve early-life organismal performance or reproduction, cause delayed epigenetic displacement, and thereby contribute to aging, while positive selection acts on the heritable genes controlling that chromatin transition.

This leads to the following restricted scale-general definition. The SAI information-theoretic framework can describe state stability and active destabilization at molecular, cellular, organismal, population, and intergenerational levels, but a particular process constitutes SAI only when heritable mechanisms influence the probability distribution of future states and the instability generated increases reproductive fitness directly or indirectly.

This qualification distinguishes a scale-general mathematical framework from a claim that biological instability is generally adaptive. Regulated state change, elevated entropy, or increased heterogeneity alone is insufficient evidence for SAI.

### 3.4. Model of passive error and active destabilization

Consider an organism containing a relatively invariant core information channel J_core_ and an environmentally responsive exploratory information system X. Let M_X_ denote maintenance investment in X. Maintenance is costly, and k_X_ M_X_ represents its immediate energetic or opportunity cost. The separation of J_core_ from X allows high fidelity maintenance and selected instability to act on different information channels rather than requiring contradictory selection on the same channel.

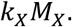

> Here J_core_ is the relatively invariant core information channel, X is the environmentally responsive exploratory information system, M_X_ is maintenance investment in X, k_X_ is the marginal energetic or opportunity cost per unit maintenance, and k_X_ M_X_ is the total immediate maintenance cost.

Passive error in the exploratory channel declines with maintenance according to

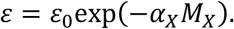

> Here ε is the passive error probability, ε_0_ is passive error in the absence of maintenance, α_X_ is the efficiency with which maintenance suppresses passive error in exploratory system X, M_X_ is maintenance investment in X, and exp denotes the exponential function.

The environment persists with probability 1 − q and changes with probability q. When the environment persists, retention of the current exploratory state is favored. When the environment changes, an alternative state becomes favorable. Conditional on passive error, let θ be the probability that the resulting alternative is appropriate in the changed environment. The organism may additionally activate a destabilization mechanism with probability a. Conditional on active destabilization, let η be the probability of generating an appropriate alternative. The focal case is

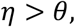

> Here η is the conditional probability that active destabilization generates an appropriate alternative after environmental change, θ is the corresponding conditional probability for passive error, and > denotes greater than.

in this way active destabilization samples recurrently useful alternatives more efficiently than passive error (Figure 1). This does not require that a mutation be produced because it will be beneficial. It requires only that selection over prior generations has shaped the statistical distribution of active variation.

**Figure 1.**
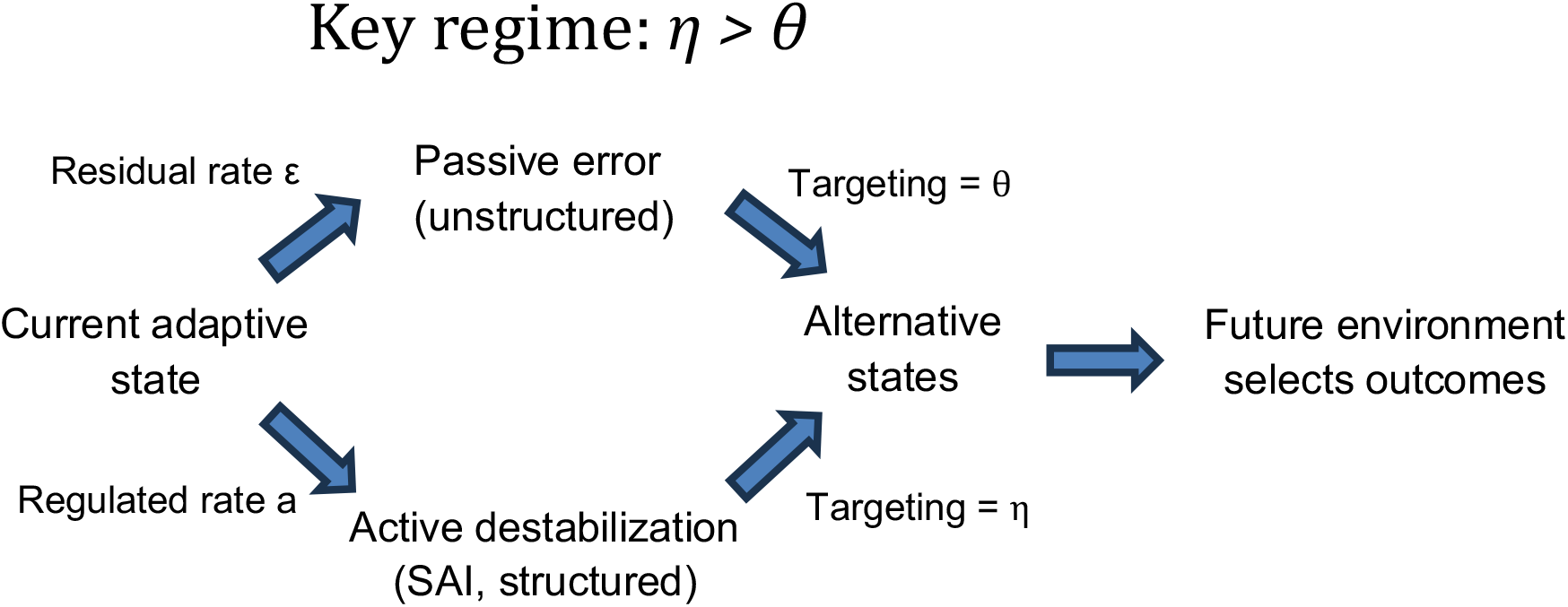
Passive versus active destabilization. Conceptual distinction between passive, comparatively unstructured error and regulated active destabilization. Active destabilization can be retained when it samples recurrently useful alternative states more efficiently than passive error (η > θ).

Let active destabilization have direct cost ca. When passive error and active destabilization occur in the same transition, the outcome is assigned to the passive route. This tie breaking convention is conservative when *η* > *θ* because the joint event is credited to the less efficient mechanism. When the environment persists, passive error is assumed to remove the system from the appropriate exploratory state. Thus accidental return to the same appropriate state is neglected, an approximation that is most appropriate when the accessible state space is large.

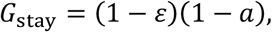

> Here G_stay_ is the probability of retaining an appropriate state when the environment persists, ε is passive error probability, a is active destabilization probability, and 1 − ε and 1 − a are the corresponding probabilities that each process does not occur.

whereas after environmental change it is

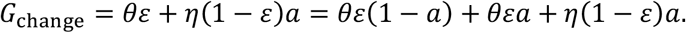

> Here G_change_ is the probability of producing an appropriate state after environmental change. The quantities θ, ε, η, and a are defined above, and the second equality expands the mutually assigned routes used by the model.

A simplifying assumption of this expected log fitness model is that each period begins with the organism matched to the current environment. G_stay_ therefore represents retention of an initially appropriate state, and mismatch is not propagated explicitly between periods. A full Markov model over joint organism and environmental states would be required when mismatch persists across periods. Passive error during a persisting environment is also assumed not to return the system accidentally to the same appropriate state, an approximation that is most appropriate when the accessible state space is large. These assumptions make the analytical model transparent but restrict its quantitative interpretation.

Ignoring fitness terms independent of ε and a, long-run log growth is

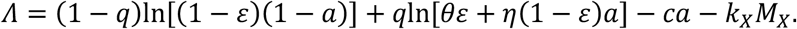

> Here Λ is long term log growth, q is the probability of environmental change, ln denotes the natural logarithm, c is the direct cost coefficient of active destabilization, and the remaining symbols are defined above. The costs ca and k_X_ M_X_ are subtracted in log fitness and therefore correspond to multiplicative fitness discounts exp(−ca) and exp(−k_X_ M_X_).

#### 3.4.1 Result 1 - active destabilization has an invasion threshold

Differentiating Λ with respect to a and evaluating at a = 0 gives

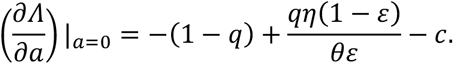

> Here ∂Λ/∂a is the partial derivative of long term log growth with respect to active destabilization probability a, and the vertical bar with a equal to zero denotes evaluation at a = 0. The remaining symbols are defined above.

Thus active destabilization can invade when

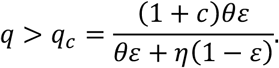

> Here q_c_ is the critical environmental change probability above which active destabilization can invade, q is the actual environmental change probability, and the remaining symbols are defined above.

Active destabilization is favored by greater environmental change q, greater targeting efficiency η, lower usefulness of passive errors θ, lower direct cost c, and lower passive error ε (Figure 2). The last relationship is important because increasing fidelity can strengthen, rather than weaken, selection for a dedicated active destabilization mechanism.

**Figure 2.**
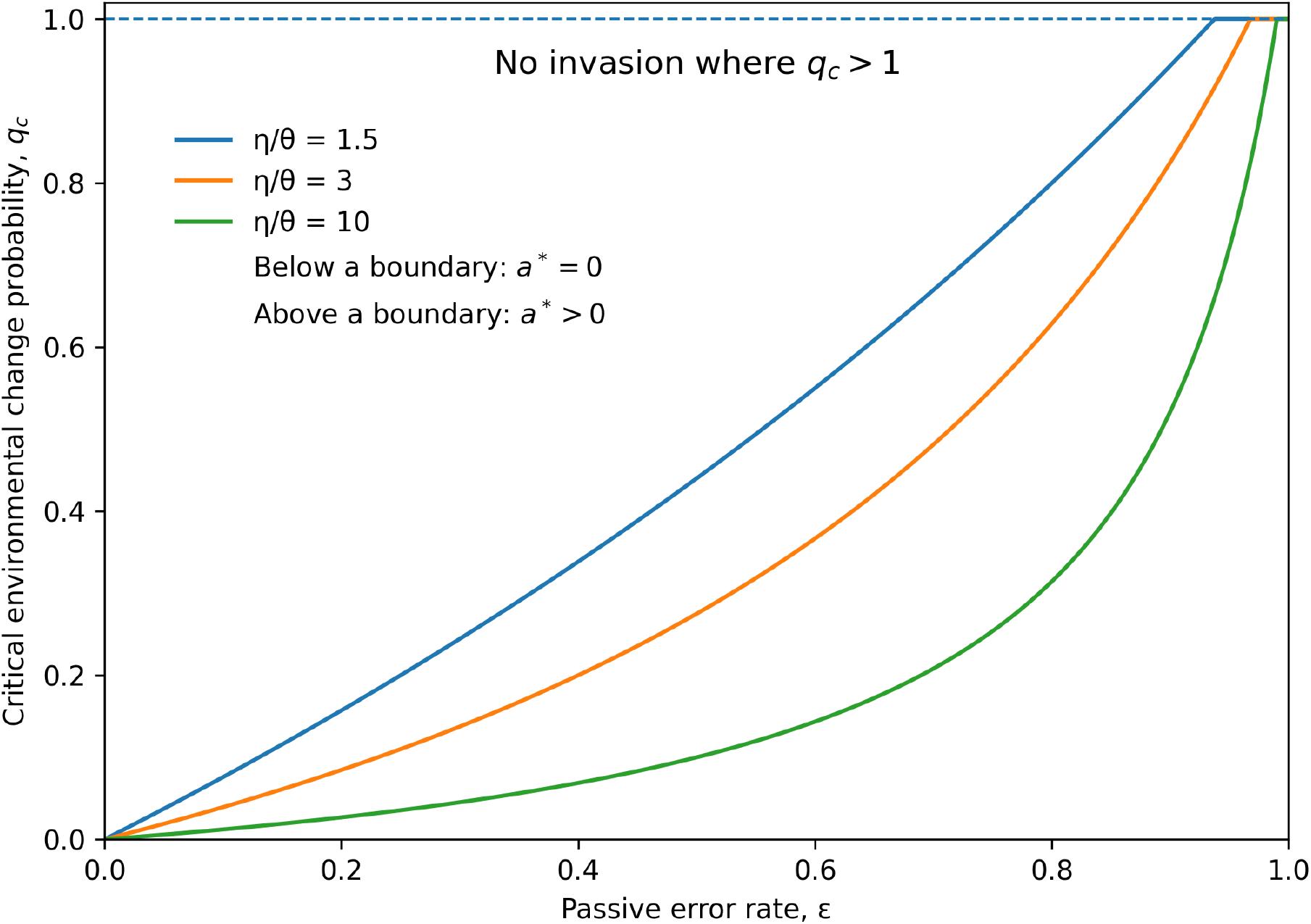
Phase boundary for invasion of active destabilization. Critical environmental change probability q_c_ as a function of passive error ε for several ratios of active to passive targeting efficiency (η/θ), with c=0.10. Above each boundary, the optimal active destabilization rate is positive. The boundary is clipped at q_c_=1 because q is a probability. Parameter combinations with q_c_>1 admit no invasion for any q∈[0,1].

#### 3.4.2 Result 2 - stabilization-destabilization complementarity is the central information-theoretic result

The interaction between active destabilization and passive error is obtained directly from the cross partial derivative

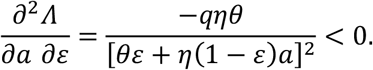

> Here ∂^2^Λ/∂a∂ε is the change in the marginal fitness effect of active destabilization a as passive error ε changes, q is environmental change probability, η is active targeting efficiency, θ is passive targeting efficiency, and < 0 denotes a strictly negative cross partial for all admissible positive parameters.

Thus reducing passive error always increases the marginal value of active destabilization. Equivalently, increasing passive error always reduces the marginal value of active destabilization. This exact result establishes stabilization destabilization complementarity without an additional sufficient condition.

For completeness, the direct marginal effect of passive error is

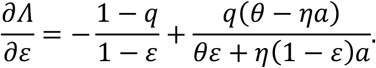

> Here ∂Λ/∂ε is the partial derivative of long term log growth with respect to passive error probability ε, and the remaining symbols are defined above.

Passive error is selected against exactly when

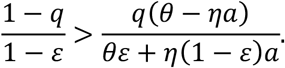

> Here the inequality is the necessary and sufficient condition for ∂*Λ*/∂*ε* < 0. The previously useful condition *ηa* > *θ* is sufficient but not necessary and is not required for the complementarity result.

Because *dε*/*dM*_*X*_ = −*α*_*X*_*ε*, costly maintenance is favored when its marginal benefit exceeds its marginal cost,

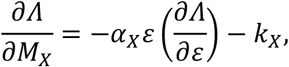

> Here ∂Λ/∂M_X_ is the marginal effect of maintenance investment M_X_ on long term log growth, and α_X_, ε, ∂Λ/∂ε, and k_X_ are defined above.

in this way an interior maintenance optimum satisfies

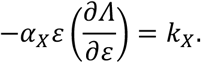

> Here the left side is the marginal fitness benefit obtained by maintenance through suppression of passive error, and k_X_ is the marginal maintenance cost. Equality defines an interior maintenance optimum.

This makes complementarity a genuine tradeoff between two costly investments. Selection can simultaneously maintain a positive active destabilization rate and invest in passive error suppression up to the point where the marginal fitness benefit of maintenance equals k_X_.

This regime is referred to as stabilization-destabilization complementarity. Selection suppresses unstructured noise while promoting structured exploration.

This result is not a statement that selection favors either low entropy or high entropy. It is a source-resolved statement about biological uncertainty. If passive error and active destabilization contribute to the same aggregate conditional entropy, selection can nevertheless drive their rates in opposite directions because the two mechanisms populate state space differently. Increasing fidelity against passive error can therefore coexist with and strengthen selection for a separate generator of structured variation.

#### 3.4.3 Result 3 - near-perfect passive fidelity can coexist with persistent active instability

In the high-maintenance limit *ε* → 0, the active-destabilization component becomes

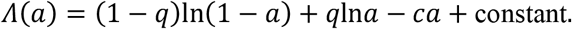

> Here Λ(a) is long term log growth as a function of active destabilization a in the high maintenance limit, constant denotes terms independent of a, and q, ln, and c are defined above.

The optimum satisfies

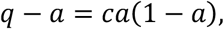

> Here q is environmental change probability, a is active destabilization probability, c is its direct cost coefficient, and the equality is the first order condition for the optimum.

giving

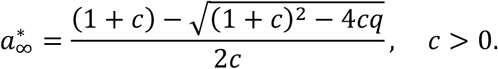

> Here a*∞ is the optimal active destabilization probability in the high maintenance limit, the asterisk denotes an optimum, the subscript ∞ denotes the limiting high maintenance solution, √ denotes the principal square root, c is the direct cost coefficient, and q is environmental change probability.

As *c* → 0, 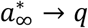. For every *q* > 0 and finite 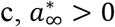 in this limiting model (Figure 3). The positivity is driven by the assumption that once passive error is eliminated (*ε* = 0), active destabilization is the only route to an appropriate state after environmental change, *G*_change_ = *ηa*, so *q*lna → −∞ as *a* → 0. Therefore the result demonstrates that near-perfect passive fidelity and selected active instability can coexist under the model’s stated transition structure, but it is not universal. If an adapted changed state remains reachable through an additional baseline route, an optimum at *a*^∗^ = 0 can become possible. The invasion threshold in Result 1 is therefore the more general statement.

**Figure 3.**
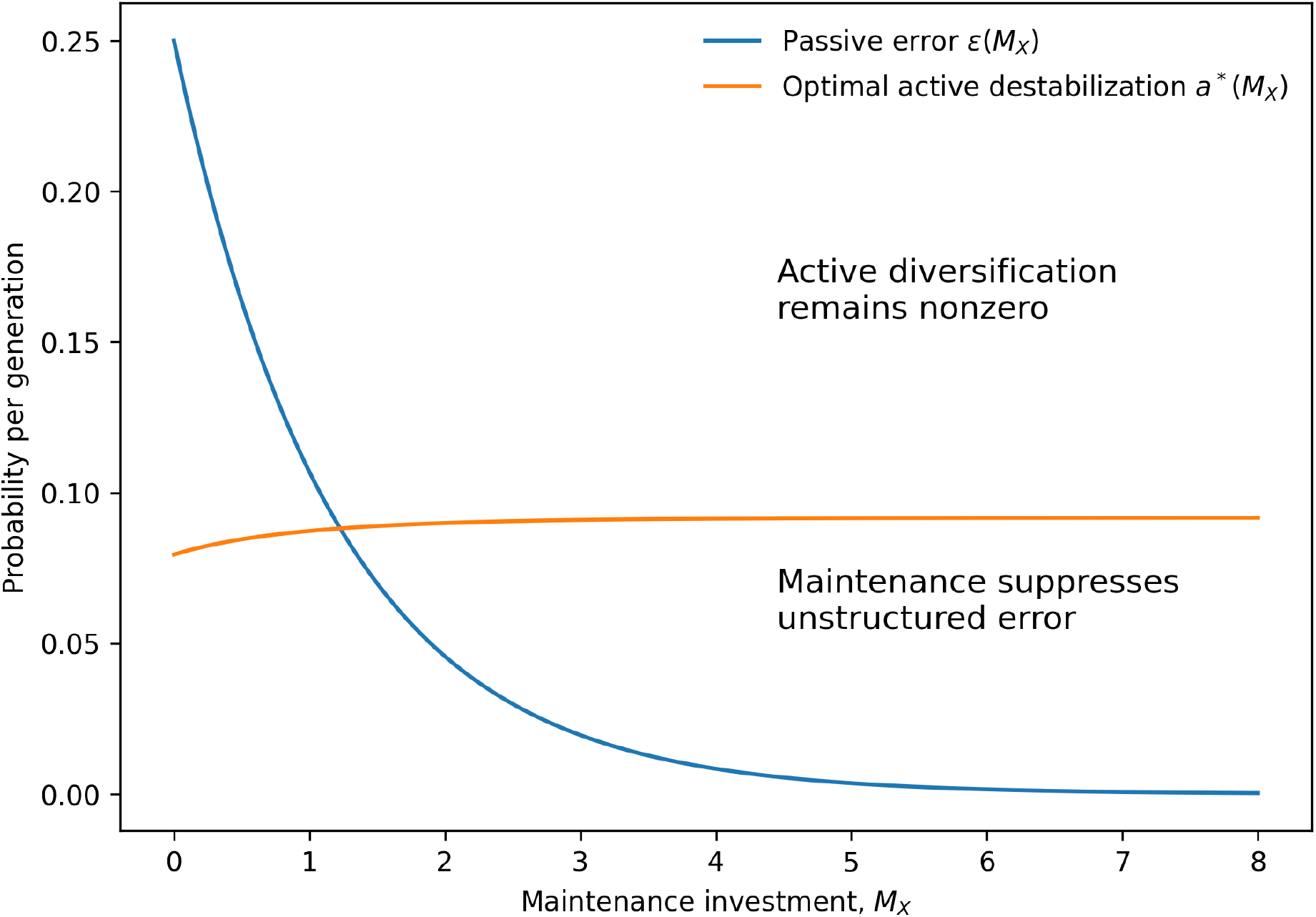
High maintenance can coexist with persistent active destabilization. Illustrative numerical solution in which increasing maintenance suppresses passive error toward zero while optimal active destabilization remains positive. Parameters are q=0.10, θ=0.02, η=0.50, c=0.10, ε_0_=0.25, and α_X_=0.85.

#### 3.4.4 Structured variation and the transition kernel

The scalar parameters θ and η summarize a more general state-space distinction. Let recurrent environmental transitions from state i be described by P_i_, passive errors by R_i_, and active diversification by A_i_. Active destabilization constitutes structured exploration when its distribution better approximates recurrent adaptive transitions, for example when

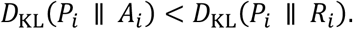

> Here D_KL_ is Kullback Leibler divergence, P_i_ is the distribution of fitness relevant future states from current state i, A_i_ is the distribution generated by active destabilization from state i, R_i_ is the distribution generated by passive error from state i, i indexes the current state, || separates the reference distribution from the comparison distribution in the divergence, and < denotes less than.

Two mechanisms can therefore generate the same amount of entropy *H*(*X*_*t*+*Δt*_ | *X*_*t*_) yet differ substantially in fitness. The relevant object is the complete conditional transition kernel, not entropy alone. The model replaces the typical scalar tradeoff “fidelity versus variation” with the distinction “unstructured error versus structured exploration.”

### 3.5 Age-structured extension and aging

A simple multiplication of all costs and benefits by the same age specific reproductive value factor cannot produce an age trend because the weighting does not alter the optimum. Temporal asymmetry is instead required. Immediate or near term benefits can be produced by selected state transitions, whereas some consequences of occupying those states can persist and impose delayed costs. The adaptive transition itself can remain fully reversible while its downstream consequences are retained. This structure is linked directly to the declining force of natural selection with age and to antagonistic pleiotropy [14,16,17].

Let x denote adult age and ω(x) the relative selective weight placed on consequences occurring after age x. Across ages where the force of selection declines, *ω*′(*x*) < 0.

#### 3.5.1 Age-dependent maintenance

Assume maintenance investment m generates future benefit 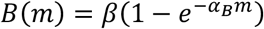 and has immediate cost k_m_ m. The age specific maintenance submodel follows the disposable soma logic that maintenance is favored according to its expected future fitness return [18].

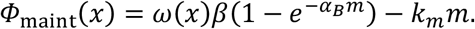

> Here Φ_maint_(x) is the age specific fitness contribution of maintenance, x is adult age, ω(x) is the relative selective weight on delayed consequences, β is the maximum future benefit of maintenance, e is the base of the natural logarithm, α_B_ controls diminishing returns of maintenance benefit, m is maintenance investment, and k_m_ is its immediate marginal cost.

The optimum is

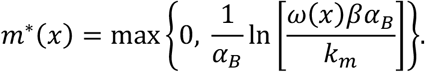

> Here m*(x) is optimal maintenance at age x, the asterisk denotes an optimum, max{·} selects the larger of the listed values and thereby constrains maintenance to be nonnegative, ln denotes the natural logarithm, and the remaining symbols are defined above.

For an interior optimum,

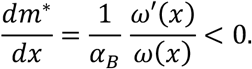

> Here dm*/dx is the derivative of optimal maintenance with respect to age, ω′(x) is the derivative of ω(x) with respect to age x, and < 0 denotes a decline with age on the interior branch. The remaining symbols are defined above.

Thus declining weight on future consequences reduces optimal maintenance on the interior branch. Once *ω*(*x*)*βα*_B_ ≤ *k*_*m*_, the nonnegativity constraint binds, *m*^∗^ = 0, and *dm*^∗^/*dx* = 0 rather than continuing to decline. The maintenance and destabilization expressions in this age extension are deliberately optimized as separate submodels rather than as a single joint age specific optimum. Their opposing trajectories illustrate the consequence of placing declining reproductive value on delayed maintenance benefits and delayed destabilization costs. A fully joint age structured allocation model would require putting both mechanisms on a common fitness scale and specifying their shared resource constraint.

#### 3.5.2 Age-dependent active destabilization

In the high fidelity limit ε approaching zero, an immediate adaptive term is produced by active destabilization, an immediate cost ca is incurred, and a delayed persistent cost c_D_a is produced by the state made accessible through destabilization. The delayed term is not required to represent failed reconstruction of the destabilized factor. It can instead represent a persistent consequence generated while the adaptive state is occupied. The effective cost is

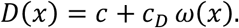

> Here D(x) is the effective age specific cost of active destabilization, c is its immediate cost coefficient, c_D_ is the coefficient for persistent delayed consequences of the explored state, and ω(x) is the relative selective weight on delayed consequences. The closed form applies for D(x) > 0. In the zero cost limit D(x) → 0, a*(x) → q.

The age-specific optimum is

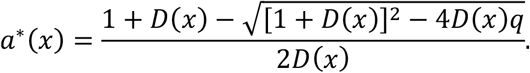

> Here a*(x) is the optimal active destabilization probability at age x, the asterisk denotes an optimum, D(x) is the effective age specific cost, q is environmental change probability, and √ denotes the principal square root.

Because ω(x) decreases with age, D(x) decreases when *c*_*D*_ > 0. Because optimal a decreases with destabilization cost,

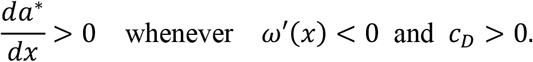

> Here da*/dx is the derivative of optimal active destabilization with respect to age, ω′(x) is the derivative of the selective weight on delayed consequences with respect to age, c_D_ is the positive coefficient of delayed cost, > 0 denotes an increase with age, and < 0 denotes a declining selective weight. If *c*_*D*_ = 0, D(x) is age invariant and *da*^∗^/*dx* = 0.

The same decline in the selective weight of delayed consequences can therefore produce opposite trajectories in which maintenance is reduced while active destabilization is retained or increased (Figure 4). A reversible exploratory transition can consequently remain favored even when repeated occupation of the resulting state produces persistent later costs.

**Figure 4.**
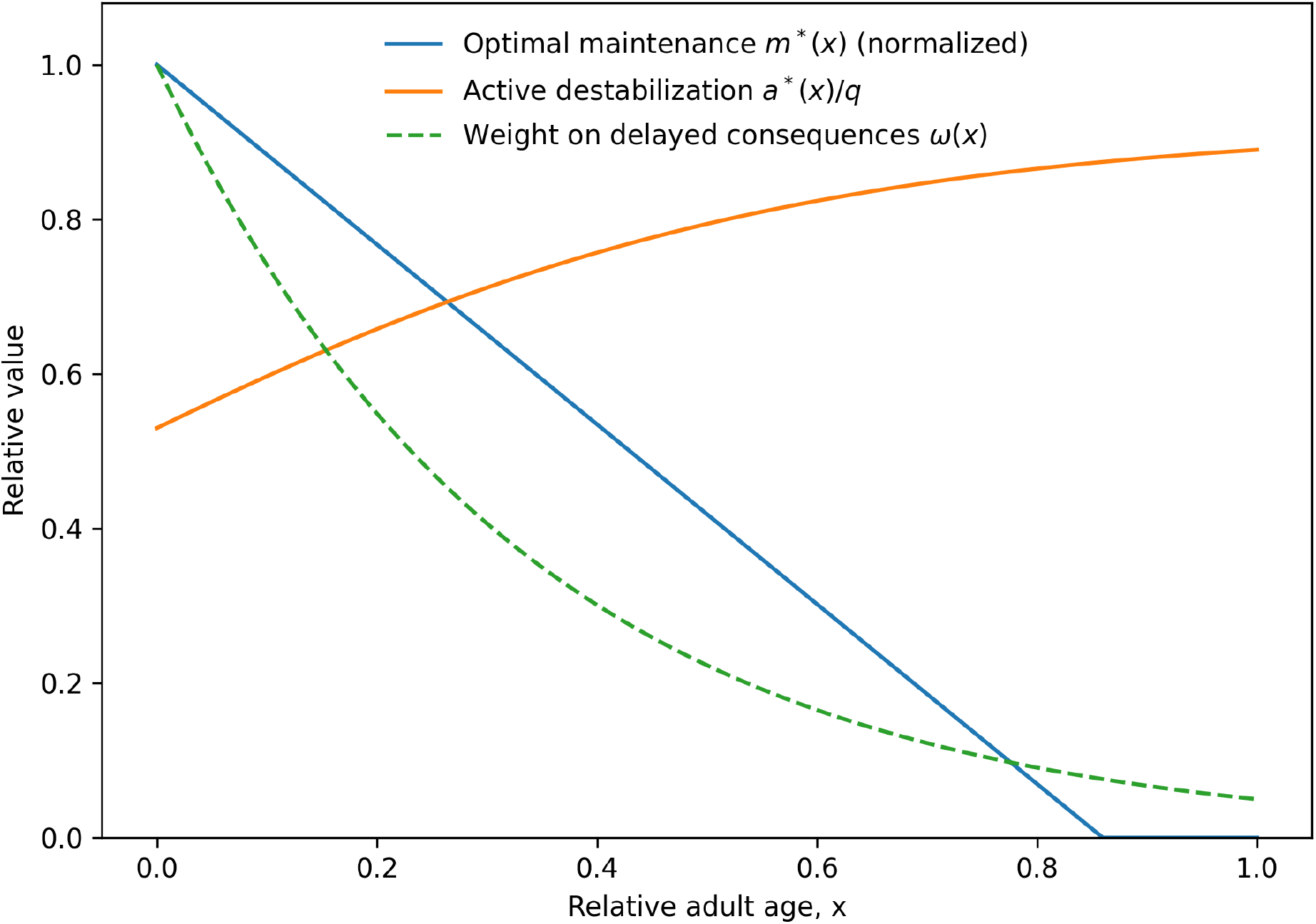
Age dependent divergence of maintenance and active destabilization. Illustrative age structured trajectories from the separate maintenance and destabilization submodels. As the selective weight on delayed consequences ω(x) declines, optimal maintenance m*(x) decreases to its nonnegative boundary while optimal active destabilization a*(x) increases toward its immediate benefit optimum. Illustrative parameters were ω(x)=exp(−3x), α_B_=1, β=1, k_m_=0.0758, q=0.10, c=0.0937, and c_D_=0.843. The maintenance curve was normalized to its value at x=0. Active destabilization is plotted as a*(x)/q, relative to the cost free environmental matching optimum approached as destabilization cost tends to zero. The selective weight ω(x) is shown on the same relative scale.

#### 3.5.3 Implications for aging and antagonistic pleiotropy

Age associated informational deterioration is divided by the model into three evolutionarily distinct categories. Unavoidable passive error is produced despite selection. Reduced maintenance is produced when future benefits receive less selective weight. Active information displacement is produced when a state changing mechanism is itself positively selected. In the third category, the exploratory transition can be fully reversible while persistent downstream consequences of the transient adaptive state accumulate.

In the aging application, SAI is interpreted as a molecular and informational realization of antagonistic pleiotropy rather than as an alternative evolutionary mechanism [3,4,6]. When a heritable mechanism increases regulated information turnover or state change, an early fitness benefit can be generated by access to an adaptive state while a delayed cost is generated by persistent consequences of occupying that state. The original factor can subsequently be restored without restoration of the complete biological state. Aging itself is not required to be adaptive. The destabilizing mechanism can be favored because of an earlier benefit while delayed deterioration is retained as a downstream consequence.

### 3.6 Sexual antagonistic pleiotropy

The fitness value of retaining versus destabilizing biological information need not be identical in females and males. Sexes can differ in reproductive schedules, endocrine signaling, energetic allocation, environmental exposure, and the fitness consequences of specific physiological states. Consequently, optimal active destabilization can be sex-specific.

Let sex *s* ∈ {*F, M*}. In the age-structured high-fidelity limit, at a specified age x, define

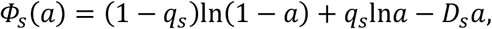

> Here Φ_s_(a) is sex specific log fitness as a function of active destabilization, s indexes sex and takes F for female or M for male, q_s_ is the sex specific environmental change or benefit probability, a is active destabilization probability, D_s_ is the sex specific effective cost, and ln denotes the natural logarithm.

Therefore

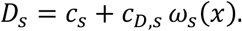

> Here D_s_ is the sex specific effective cost evaluated at the specified age x, c_s_ is the sex specific immediate cost coefficient, c_D,s_ is the sex specific delayed cost coefficient, ω_s_(x) is the sex and age specific selective weight on delayed effects, and x is age. The sex specific closed form applies for Ds > 0. In the zero cost limit Ds → 0, a*s → qs.

If each sex could evolve independently,

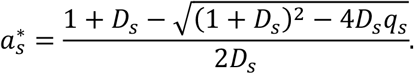

> Here a*_s_ is the sex specific optimal active destabilization probability, the asterisk denotes an optimum, and D_s_, q_s_, and √ are defined above.

In general, 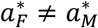. The difference 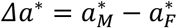 measures divergence between the sex-specific informational optima.

#### 3.6.1 Shared genetic control and intralocus sexual conflict

Let a single genetic modifier g regulate active destabilization in both sexes, such that *a*_*F*_ = *a*_*M*_ = *g*. The sex-specific selection gradient is

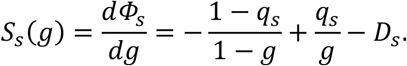

> Here g is a shared genetic modifier controlling active destabilization in both sexes, S_s_(g) is the sex specific selection gradient on g, dΦ_s_/dg is the derivative of sex specific log fitness with respect to g, and q_s_ and D_s_ are defined above.

Each sex-specific fitness function is strictly concave because

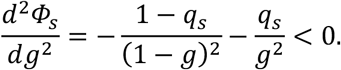

> Here d^2^Φ_s_/dg^2^ is the second derivative of sex specific log fitness with respect to g, and < 0 denotes strict concavity. The remaining symbols are defined above.

Assume a*_F_ < a*_M_. Then any shared value satisfying

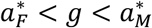

> Here a*_F_ and a*_M_ are the female and male optimal active destabilization probabilities, respectively, g is the shared modifier value, and the inequalities specify a shared value lying between the two sex specific optima.

produces *S*_*F*_(*g*) < 0 but *S*_*M*_(*g*) > 0. An allele increasing g is therefore selected against in females and favored in males. This is sexual antagonistic pleiotropy expressed through information stability.

#### 3.6.2 Population compromise and sex-specific load

Let w_F_ and w_M_ denote the relative contributions of female and male fitness to selection, with *w*_*F*_ + *w*_*M*_ = 1. Population log fitness is

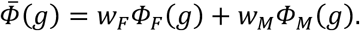

> Here 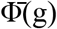 is population mean log fitness at shared modifier value g, w_F_ and w_M_ are the relative female and male contributions to selection, and Φ_F_(g) and Φ_M_(g) are female and male log fitness.

For this model, define 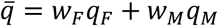 and 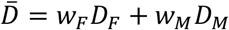. Because *w*_*F*_ + *w*_*M*_ = 1, 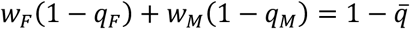, so the weighted fitness 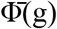 collapses exactly, not approximately, to the same single-sex functional form with parameters 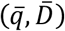. The shared optimum is therefore exactly

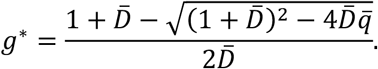

> Here g* is the exact population optimal shared modifier, the asterisk denotes an optimum, 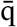 is the sex weighted mean environmental change or benefit probability, 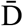 is the sex weighted mean effective cost, and √ denotes the principal square root. This closed form applies for 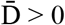. In the zero cost limit 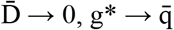.

When both sexes contribute positively to selection and their optima differ, g* lies between them. Neither sex reaches its own optimum. Define the information-stability sex-specific load as

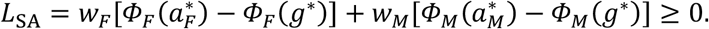

> Here L_SA_ is the sexual antagonism load, w_F_ and w_M_ are the relative female and male contributions to selection, Φ_F_ and Φ_M_ are sex specific log fitness functions, a*_F_ and a*_M_ are their separate optima, g* is the shared population optimum, and ≥ 0 denotes a nonnegative load.

Near the optima, write 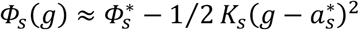, where the local curvature is

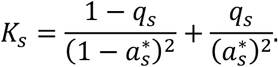

> Here K_s_ is the magnitude of local curvature of sex specific log fitness at its optimum and therefore measures local stabilizing selection, q_s_ is the sex specific environmental change or benefit probability, and a*_s_ is the sex specific optimum.

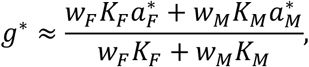

> Here K_F_ and K_M_ are the female and male local curvature values, ≈ denotes the local quadratic approximation, and the remaining symbols are defined above.

and the sex-specific load becomes

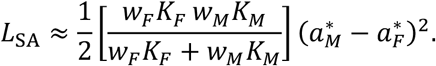

> Here ½ is one half, L_SA_ is the sexual antagonism load, and the remaining symbols are defined above. The expression is the local quadratic approximation to the exact load.

Therefore, to first approximation, unresolved sexual conflict load increases with the square of the difference between male and female optimal destabilization rates (Figure 5). Both the curvature weighted approximation for g* and the quadratic approximation for L_SA_ are local approximations. The curvature weighted g* is presented for intuition only because the exact closed form is available and is used elsewhere. Numerical checks show that the approximations deteriorate as 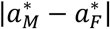 increases. For the Figure 5 parameter set, where 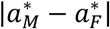 is approximately 0.103, the quadratic load approximation underestimates the exact load by approximately 15 percent and the curvature weighted g* approximation underestimates the exact g* by approximately 33 percent.

**Figure 5.**
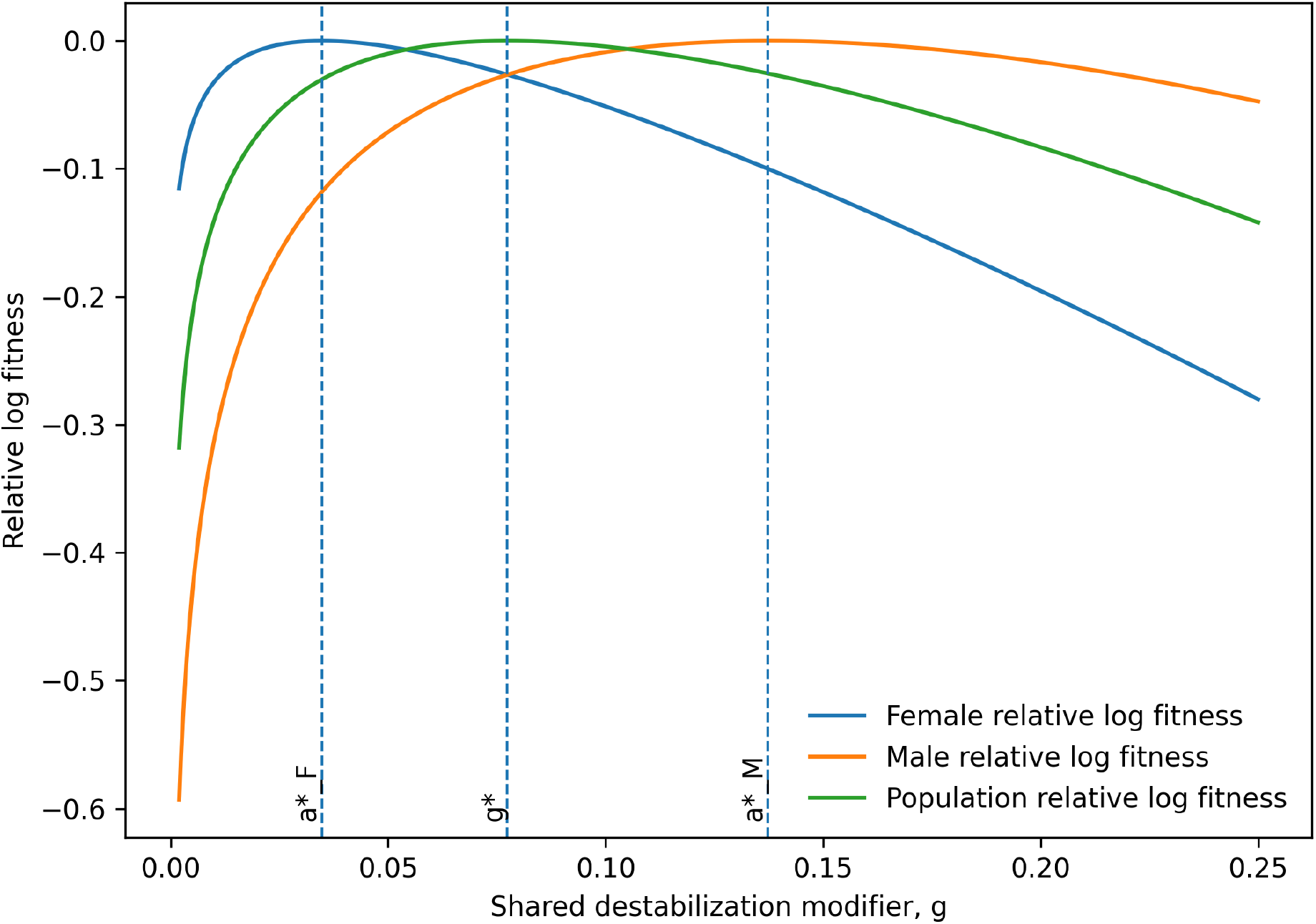
Shared regulation creates sexual conflict over information stability. Illustrative female and male log-fitness surfaces for a shared destabilization modifier g. Different sex-specific optima force the population optimum g* to lie between a*_F_ and a*_M_, generating unresolved intralocus sexual conflict and a positive sex-specific load. Illustrative parameters were q_F_=0.06, D_F_=0.75, q_M_=0.18, D_M_=0.36, w_F_=0.52, and w_M_=0.48. These values give a*_F_=0.0348, g*=0.0774, and a*_M_=0.1373. Curves are displayed as relative log fitness after subtracting each curve’s maximum for visualization. Both q_F_ < q_M_ and D_F_ > D_M_ contribute to the illustrated ordering of the sex specific optima, so the figure is not intended to isolate either mechanism.

#### 3.6.3 Sex-specific regulation and resolution of conflict

A second modifier that permits independent regulation, *g* → (*g*_*F*_, *g*_*M*_), can allow each sex to approach its own optimum. Regulatory decoupling is favored when the recoverable sex-specific load exceeds the cost of maintaining sex-specific control. The framework therefore predicts stronger selection for sex-biased regulation when divergence between male and female information-stability optima is large.

The age and sex extensions interact naturally because *D*_*s*_(*x*) = *c*_*s*_ + *c*_*D,s*_ *ω*_*s*_(*x*). If delayed costs lose selective importance at different rates in the sexes, *Δa*^∗^(*x*) can itself change with age. Sexual conflict over information stability can therefore increase or decrease across the reproductive lifespan. A single allele may simultaneously exhibit age dependent antagonistic pleiotropy and sexual antagonistic pleiotropy.

#### 3.6.4 Implications for sexual antagonistic pleiotropy

The same distinction applies to sex-specific optima. Females and males may differ both in the optimal amount of transition entropy and in the optimal transition kernel. Let H_s_* denote the sex specific optimal transition entropy and Q_s_* the sex specific optimal transition kernel. Thus 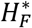 need not equal 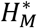, and more importantly 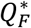 need not equal 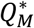. Shared genetic regulation can force a compromise kernel that is suboptimal in both sexes, generating intralocus sexual conflict and the sex-specific load derived above. Sexual conflict can therefore concern not merely how much biological uncertainty is generated, but where that uncertainty is directed in state space [19–21].

## 4. Discussion

### 4.1 SAI and Shannon information theory

Information theory has found broad applications in biology, including systems biology, genomics, metabolomics, neuroscience and evolution [22–25]. These studies have been facilitated by the development of artificial intelligence and large language models (LLMs), which have transformed several areas of biological research, including protein structure, genomics and drug discovery [26,27]. Here, the computational power of LLMs was leveraged to help derive several key equations that integrate SAI with modern information theory.

The principal result is that biological stability and selected instability can be complementary. Passive error can be strongly suppressed while a separate state-generating process is retained because recurrently useful alternatives are sampled. The relevant evolutionary object is therefore the transition kernel rather than Shannon entropy alone.

The mechanistic source, state space distribution, and temporal structure of uncertainty are more informative than total entropy alone. Conditional entropy quantifies unpredictability generated by a transition process, whereas temporal mutual information quantifies retention of information about previous states. Relative entropy quantifies mismatch between generated states and fitness relevant future states. Equal stationary state entropy can therefore be associated with different rates of forgetting and different fitness consequences.

A direct bridge between SAI and Shannon state entropy is provided by the minimal AB replicator model. Loss of B generates nonreplicating A but also permits formation of the superior AAB state. When positive B instability is selected, occupancy is necessarily spread beyond AB and Shannon state entropy is increased relative to the perfectly stable reference. Entropy is produced as a consequence of selected exploration rather than as a quantity that is itself maximized.

A more direct cellular interpretation is provided by the reversible factor turnover model. B present and B absent cellular states are generated by degradation and resynthesis of B, and a combinatorial state space can be generated by n reversible factors. Information about an earlier factor configuration can be erased by physical degradation even when the same coarse grained factor state is later restored by resynthesis. Stationary state entropy can remain unchanged while the rate at which temporal mutual information is lost is changed by proportional changes in degradation and synthesis. Selected turnover can therefore be interpreted as controlled biological forgetting when access to states suited to future environments is improved by reduced dependence on past states. Aging is not implied by reversible forgetting itself. Aging can be produced when transient adaptive states generate persistent downstream consequences that remain after the initiating factor has been restored. Imperfect reconstruction is one possible source of persistence, but it is not required.

A new Shannon theorem is not proposed. Instead, an evolutionary application is derived in which the source of uncertainty is made explicit and selected molecular instability is linked to structured exploration of biological state space. This distinction allows increased uncertainty to be adaptive even while passive error is suppressed.

### 4.2 SAI and antagonistic pleiotropy

The extension of the SAI information theory model to aging is directly consistent with antagonistic pleiotropy. Immediate or early benefits can be produced by selected state space exploration and controlled forgetting when obsolete states are discarded and alternative states improve performance under current or changing environments. Delayed costs can be produced when the transient adaptive states generate persistent downstream consequences. The initiating switch can remain fully reversible while the larger cellular state is progressively displaced. When both effects are produced by the same heritable destabilizing mechanism, antagonistic pleiotropy is generated. This relationship is not regarded as a distinct evolutionary principle introduced by SAI. Selection for molecular instability through antagonistic pleiotropy was previously modeled for a hypothetical two-subunit replicator, where increased proliferation was produced under defined conditions by instability of one subunit [6]. In the present framework, that idea is generalized by information theory across biological scales and by explicit separation of passive error from regulated destabilization. A distinct prediction is thereby produced. Retention of past factor state information should be increased by suppression of selected turnover under appropriate conditions, but adaptive plasticity or fitness under changing environments can be reduced at the same time.

### 4.3 SAI complements the ITOA

A comparison with the previously-described Information Theory of Aging (ITOA) [28–31] is especially useful, because different levels of the same potentially causal sequence can be represented by the two frameworks. In the ITOA, aging has been proposed to be driven by progressive loss of youthful epigenetic information [28]. Relatively stable genetic information is distinguished from more environmentally responsive epigenetic information in this framework, and retrieval of youthful states by epigenetic reprogramming is emphasized. A direct physical connection to the present SAI model can be made because epigenetic chromatin states are implemented by distributions, modifications, interactions, turnover and localization of molecular factors. Chromatin components and regulators can therefore be represented among the factors F_1_ through F_n_ whose turnover and redistribution permit movement among cellular states, as described above.

A causal role for epigenetic information displacement in aging was supported by experiments in which repeated induction and faithful repair of DNA breaks accelerated molecular and physiological features of aging without a corresponding mutational burden [31]. The repair response should not, however, be described as random or unstructured. Regulated redistribution of chromatin modifying proteins during DNA damage responses had previously been shown to promote DNA repair and genomic stability while also producing transcriptional changes resembling those observed during aging [32]. In this way a biologically beneficial and structured response can have a persistent informational consequence.

The distinction proposed here is therefore evolutionary rather than a distinction between structured and random information loss. In ITOA, loss or displacement of youthful epigenetic information is treated primarily as a proximate cause of aging. In SAI, an additional question is asked concerning why an information destabilizing process capable of producing such displacement can be maintained or favored by natural selection. Selection can be favored when a regulated state changing process provides an earlier benefit, while a persistent informational cost can be expressed later. Importantly, the chromatin alterations associated with aging need not themselves have been directly selected. Selection is proposed to act on the regulated process that generates useful state transitions. Persistent epigenetic displacement can then arise as a delayed consequence of repeated operation of that process. The early benefit was not measured as reproductive fitness in the ICE experiments, and those experiments should therefore not be interpreted as a complete demonstration of SAI. Rather, the results are consistent with a sequence in which a selected repair response provides an immediate functional benefit and repeated operation contributes to later displacement from a youthful epigenetic state.

### 4.4 SAI and sexual antagonistic pleiotropy

If females and males have different optimal destabilization rates but share genetic control, intralocus sexual conflict follows directly. The resulting sex-specific load depends on the divergence between the sex-specific optima and can itself vary with age. This provides an information-stability interpretation of sexual antagonistic pleiotropy and predicts that sex-biased aging can arise when shared mechanisms of biological state change are differently optimized in the two sexes.

Information theoretic approaches have previously been applied to sexual selection and nonrandom mating, including the use of relative entropy to quantify information gained through nonrandom mating [33]. However, sexual conflict has generally been formulated in terms of divergent sex specific phenotypic fitness optima and shared genetic constraints [19,34–38]. The present model applies this general optimization structure specifically to sex specific biological information dynamics. It shows that intralocus sexual conflict can arise when females and males have different optimal rates or transition kernels for a regulated information destabilizing process but share genetic regulation of that process.

### 4.5 Biological interpretations and candidate systems

The SAI information theory model is intended as a general evolutionary framework rather than as a claim that all regulated variation has the same origin. Candidate systems are expected to combine strong suppression of background error with dedicated mechanisms that generate structured variation. Phenotype switching, stress induced mutagenesis, immune diversification, recombination, and regulated molecular turnover provide candidate examples. The critical empirical test is whether the actively generated transition distribution is more useful than the distribution produced by passive errors and whether the destabilizing mechanism itself carries a selectable cost.

For aging, candidate mechanisms would be regulated pathways that repeatedly remodel chromatin, transcriptional state, metabolism, tissue composition, or endocrine signaling in ways that improve early-life reproductive fitness but create persistent later information displacement. These mechanisms may be related to the hallmarks of aging and to pathways that exhibit hyperfunction [39–43]. The model is consistent with the idea that suppressing such pathways can extend late-life maintenance while decreasing the earlier function for which the pathway was selected.

For sexual antagonistic pleiotropy, the candidates are shared pathways for state change that have different reproductive value in females and males. Sex-specific hormones and chromatin regulators are obvious potential mechanisms for partial resolution of conflict [35], because they can decouple a shared genetic architecture into sex-biased destabilization phenotypes.

### 4.6 Limitations of the study

The study includes several simplifications designed to make the analytical conditions transparent. The strong-selection model compresses complex fitness landscapes into matching probabilities θ and η. Maintenance and active destabilization are represented by simple cost functions. Within each environmental regime, the probability of environmental change is assumed to be constant over time, although different regimes can have different probabilities of change. And finally sex-specific fitness is combined with fixed weights. Future models should incorporate finite mismatch fitness, variable probability of environmental change with time, explicit transition matrices, density dependence, and age-structured demographic schedules derived directly from survival and fertility functions.

### 4.7 Previous experimental support and future directions

Previous studies provide support for certain components of the model. Specifically, they distinguish selectively advantageous active destabilization from relaxed maintenance or unavoidable damage. Acar et al. engineered *Saccharomyces cerevisiae* populations that stochastically switch between two phenotypic states at experimentally tunable rates [44]. In their system, the experimentally controlled phenotypic switching produced a fitness advantage under rapid environmental change but a disadvantage under slow environmental change, showing that regulated instability can be beneficial in this engineered system in an environment-dependent manner. Abreu et al. measured fitness effects of adaptive *Saccharomyces cerevisiae* mutants in static and fluctuating environments, and showed that fitness in one component of a fluctuating environment can depend strongly on the preceding component [45]. Their analysis shows that transition history contains substantial information about fitness, supporting the model’s treatment of biological change as a structured, history-dependent information process rather than undirected loss of fidelity. These results therefore provide empirical support for the central distinction between passive error and selectively advantageous active variation.

A stronger test of the extended evolutionary model would demonstrate the complete predicted joint signature within a single biological system. For example, low background error, regulated active variation, a direct reproductive fitness benefit of that variation under specified conditions, a cost of generating it, and delayed informational or fitness consequences. For the aging and sexual-antagonism extensions specifically, the strongest tests would further demonstrate that these delayed consequences vary with age and/or sex. Such experiments would test whether an actively destabilizing mechanism that is favored because of its immediate or early-life benefits can subsequently contribute to age-dependent information loss or sexually antagonistic fitness effects.

## 5. Conclusions

Natural selection need not maximize biological information fidelity or minimize Shannon entropy. It can favor accurate preservation of existing information while separately favoring costly mechanisms that generate structured departures from that information. The central result, stabilization-destabilization complementarity, shows that passive error can be driven toward zero while optimal active destabilization remains positive. Therefore the evolutionary value of biological uncertainty depends on its source and state-space distribution, not only on its magnitude. When delayed consequences lose selective weight with age, maintenance can decline while active destabilization persists or increases. When male and female optima differ under shared genetic control, the same mechanism produces sexual antagonistic pleiotropy and a measurable sex-specific load.

The general hypothesis is that biological information is stabilized according to its predictive value for future fitness, whereas active destabilization is favored when structured exploration of alternatives has greater expected value than preservation of the current state. Standard Shannon quantities are used to formalize this evolutionary distinction. SAI of information can be expressed as antagonistic pleiotropy when an earlier benefit is accompanied by a delayed informational cost. Taken together, the results provide a unifying framework for SAI, information theory and antagonistic pleiotropy.

## Supplementary materials

Supplementary materials and methods; Supplementary Table S1: Principal symbols used in the models.

## Data Availability Statement

The code required to reproduce the reported analyses and figures are included in the accompanying reproducibility (SAI_reproducibility_archive_V23.zip). A README file, software requirements, and a manuscript to code output manifest are also included. The archive is deposited in Zenodo (<u>10.5281/zenodo.22179057</u>).

## Supplementary materials

### 1. Supplementary materials and methods

#### 1.1 Minimal AB replicator analysis

The minimal replicator state space contained AB, A, and AAB. Replication was assigned to AB and AAB, with R_A_ set to zero and R_AAB_ greater than R_AB_. Differential instability was represented by loss of B from AB at rate d_B_. Return from A to AB was represented by γ and loss from AAB was represented by u. Access to AAB was represented by χ(d_B_), with the small d_B_ expansion χ(d_B_) = k_1_ d_B_ + O(d_B_^2^). First order state occupancies and the corresponding derivatives of replication and function were obtained by expansion around d_B_ = 0. Shannon state entropy was calculated over the three state distribution. Divergence from the youthful reference distribution P_0_ = (1,0,0) was evaluated with Jensen Shannon divergence.

#### 1.2 Reversible factor turnover model

A cellular state variable was defined by the presence or absence of factor B. Degradation and synthesis rates were denoted k_d_ and k_s_. The stationary probabilities were calculated as p(B present) = k_s_ divided by k_s_ plus k_d_ and p(B absent) = k_d_ divided by k_s_ plus k_d_. Binary Shannon entropy was calculated from these probabilities. Temporal dependence was evaluated from the two state continuous time Markov process. The characteristic correlation time was calculated as 1 divided by k_d_ plus k_s_, and temporal mutual information was used to represent retention of information about an earlier B state. The model was generalized to n binary factors, giving at most 2^n^ accessible states. Environmental matching was represented conceptually through comparison of the cellular transition kernel Q with the distribution of cellular states favored under recurrent environmental transitions.

#### 1.3 Mathematical model and analytical calculations

The model was analyzed using the definitions and equations presented in Results. Long term log growth was optimized with respect to active destabilization a and, where applicable, maintenance investment M_X_. The invasion threshold was obtained from the sign of the derivative of Λ with respect to a at a=0. Passive error was modeled as ε=ε_0_ exp(−α_X_ M_X_). Stabilization destabilization complementarity was established from the exact negative cross partial ∂^2^Λ/∂a∂ε. Interior optima were obtained analytically and constrained to admissible probability or nonnegative investment domains.

#### 1.4 Information theoretic calculations

Shannon state entropy in the minimal replicator was calculated with base 2 logarithms and is therefore reported in bits. The Jensen Shannon divergence in the minimal replicator was likewise calculated with base 2 logarithms and is reported in bits, so its small-displacement limit is δ/2. Expected log fitness and standalone Kullback Leibler expressions use natural logarithms and are therefore measured in nats where an information unit is applicable. In the reversible factor turnover model, stationary state probabilities and the correlation time were derived analytically.

Temporal mutual information was used conceptually to distinguish stationary state diversity from loss of information about earlier states. No unreported numerical conditional entropy, entropy rate, temporal mutual information, or Kullback Leibler analysis is claimed.

#### 1.5 Numerical evaluation and theoretical figures

Numerical calculations were used to evaluate closed form expressions and generate the computational figures. Figure 2 evaluated q_c_(ε) with c=0.10 for η/θ ratios of 1.5, 3, and 10, with q_c_ clipped at 1 because q is a probability. Figure 3 used q=0.10, θ=0.02, η=0.50, c=0.10, ε_0_=0.25, and α_X_=0.85, with a*(ε) obtained from the admissible closed form optimum of Λ at each maintenance value. Figure 4 used ω(x)=exp(−3x), α_B_=1, β=1, k_m_=0.0758, q=0.10, c=0.0937, and c_D_=0.843. For visualization, m*(x) was normalized to m*(0), a*(x) was divided by q to express active destabilization relative to the cost free environmental matching optimum, and ω(x) was plotted directly as a relative selective weight. Figure 5 used the sex specific parameters reported in its legend. Figures 1 through 5 and the associated numerical output tables are regenerated by the master reproducibility script reproduce_all.py.

#### 1.6 Reproducibility

The theoretical figures are regenerated from the equations and parameter values reported in the manuscript by the master script reproduce_all.py. Software requirements and a manuscript to code output manifest are included in the archive.

**Supplementary Table S1.**
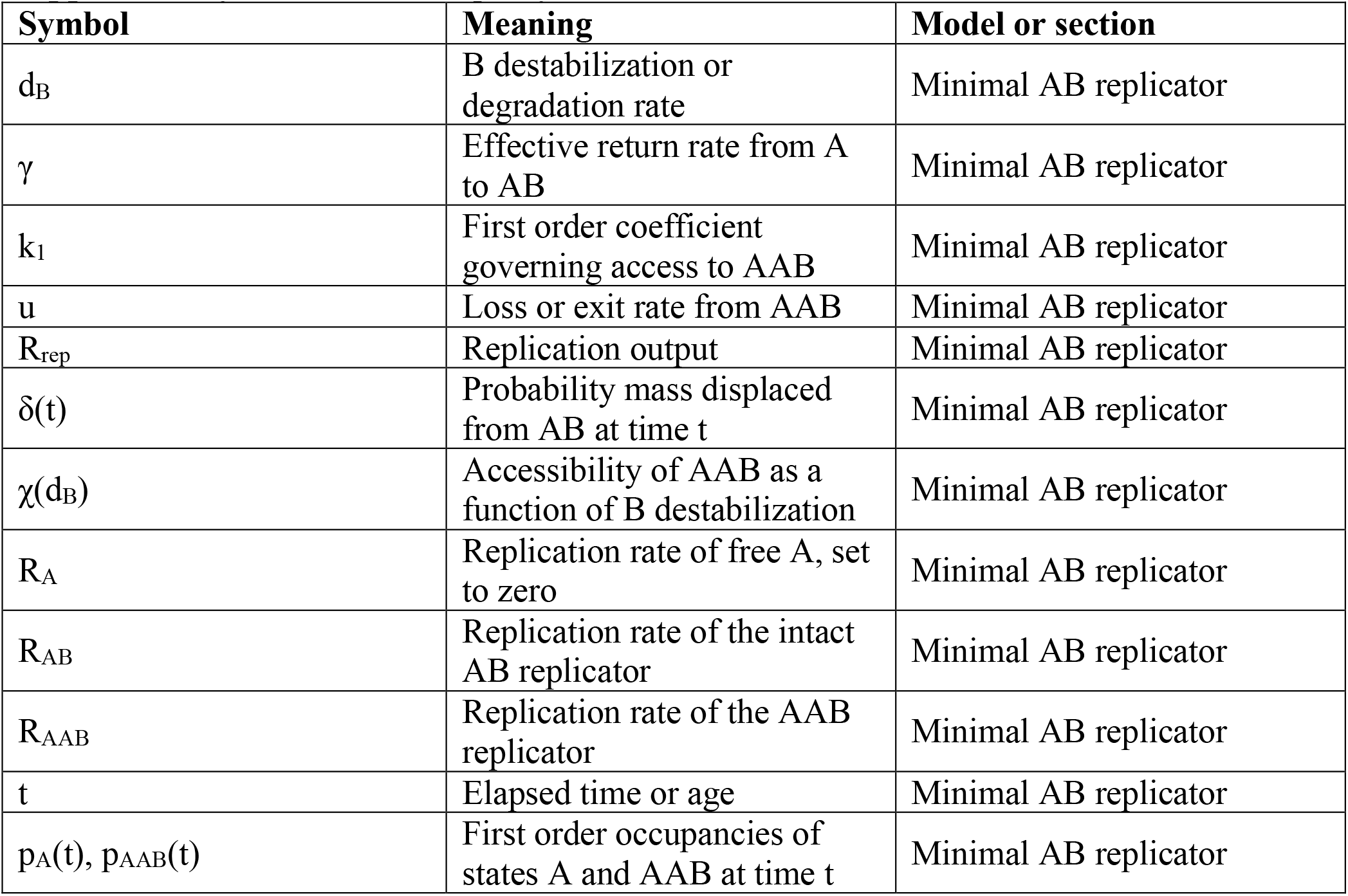
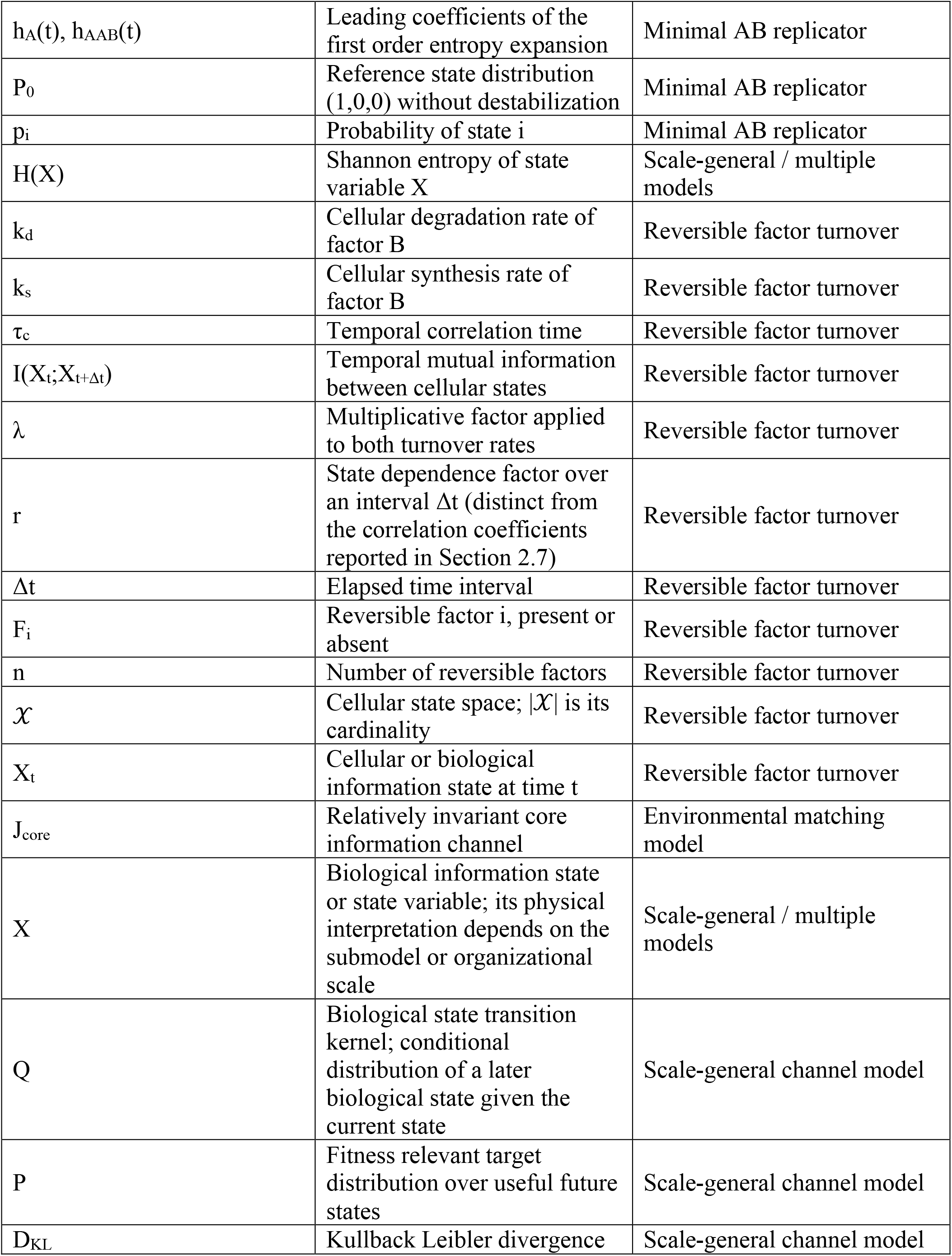
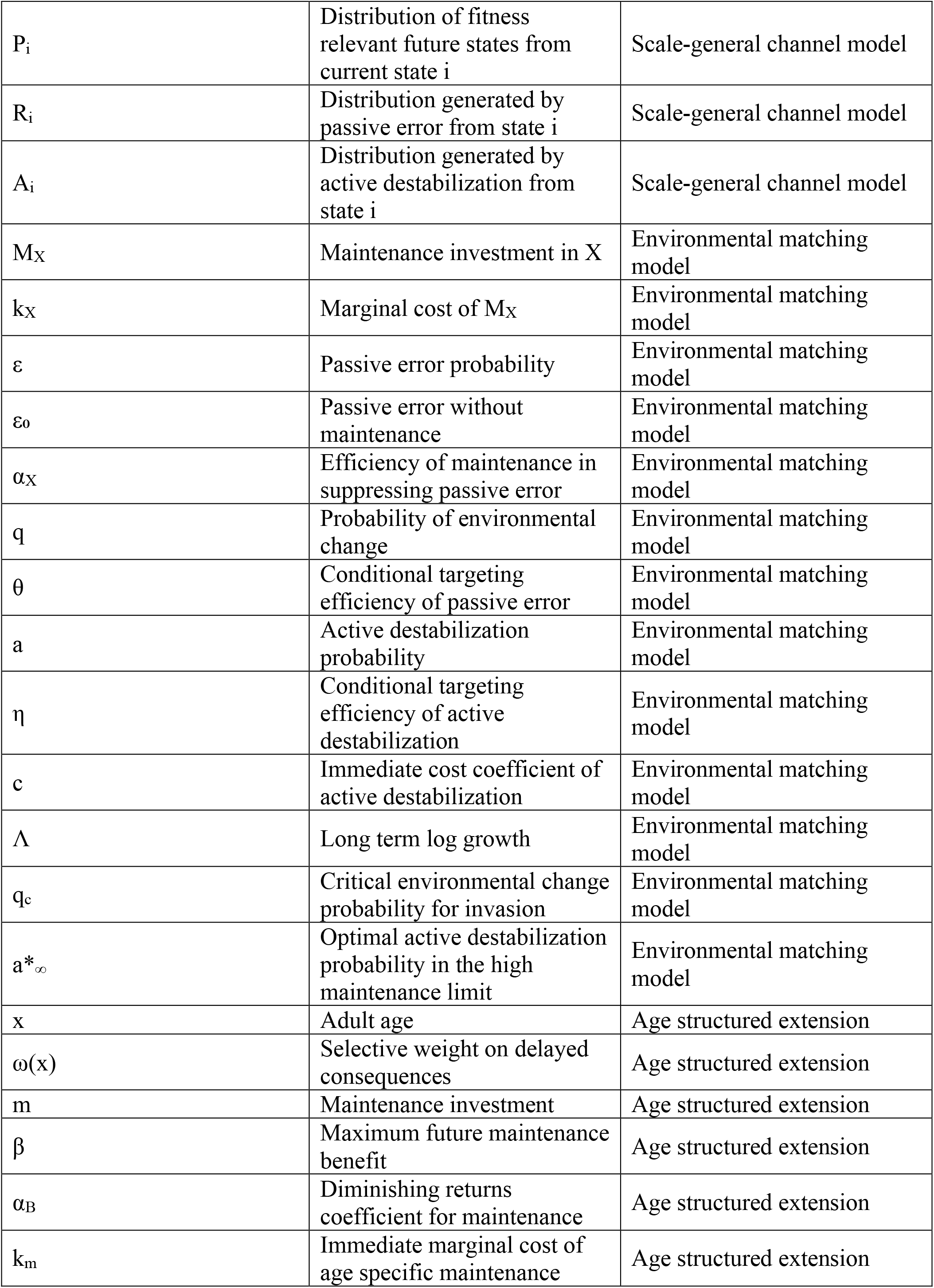
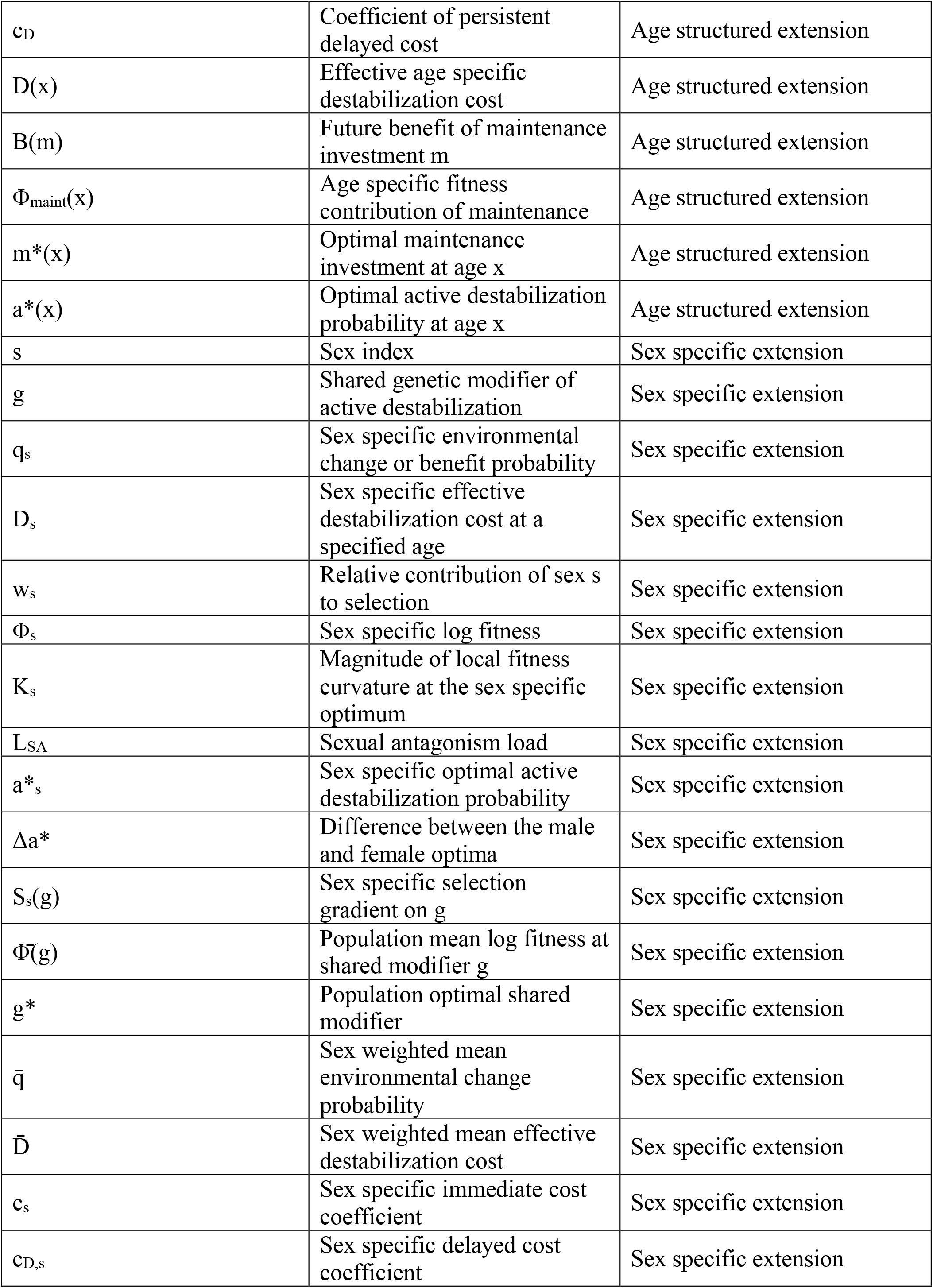
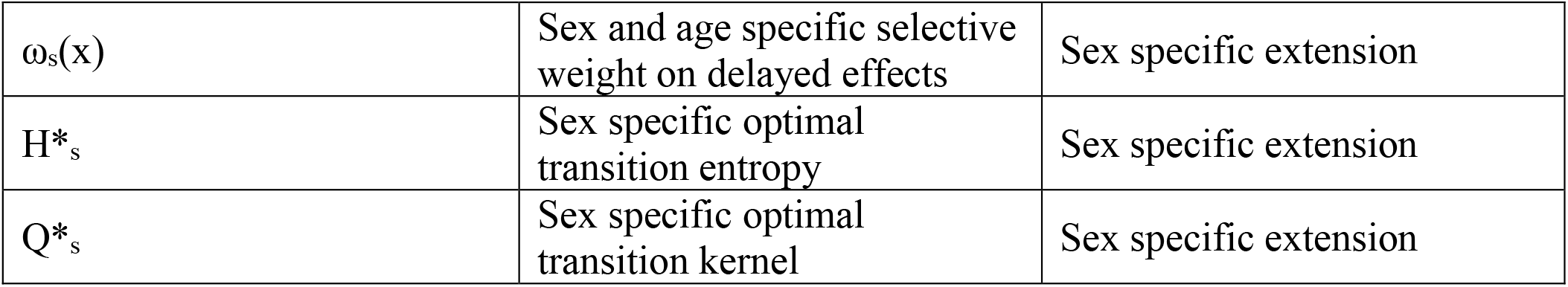
Principal symbols used in the models.

